# A matter of availability: Neural representations of task-relevant stimulus features are sharper when stimuli are memorized rather than perceived

**DOI:** 10.1101/2022.09.01.506184

**Authors:** Samson Chota, Surya Gayet, J. Leon Kenemans, Chris N.L. Olivers, Stefan Van der Stigchel

## Abstract

Our visual environment is relatively stable over time and an optimized visual system ought to capitalize on this by not devoting any representational resources to objects that are still present. Subjective experience, however, suggests that externally available (i.e., perceived) information is more strongly represented in neural signals than memorized information. To distinguish between these possibilities, we use EEG multivariate pattern analysis to quantify the strength of representation of task-relevant features (color or spatial frequency) in anticipation of a change-detection task. Perceptual availability was manipulated between experimental blocks by either keeping the stimulus on the screen during a two second delay period (perception) or removing it shortly after its initial presentation for the same time period (memory). We find that task-relevant (i.e., attended) memorized features are more strongly represented than irrelevant features. More importantly, we find significantly weaker representations for available (perceived and attended) features than for unavailable (memorized and attended) features. Contrary to what subjective experience suggests, our findings demonstrate that vividly perceived and attended stimuli elicit weaker neural representations (in terms of detectable multivariate information) than stimuli maintained in visual working memory. We hypothesize that an efficient visual system spends little of its limited resources on the internal representation of information that is externally available anyway.

## Introduction

The brain is the most energy demanding organ in the human body, consuming approximately 20% of total energy (Magistretti & Allaman, 2015). Important cognitive processes implemented by neural activity are metabolically costly and therefore need to be highly optimized. One of these crucial processes is working memory, often defined as a short term storage for information that is no longer available (Baddeley, 1992). Working memory is surprisingly limited, with the average person only being able to remember around 3-5 objects depending on their complexity and the task context (Cowan, 2010). This limitation is often explained by a resource bottleneck, further emphasizing the need for efficient distribution of resources in the brain (Cowan, 2010). Furthermore, in more ecological contexts, e.g. when giving participants the choice on how many items to encode in WM at a time, very little information is actually maintained internally (Draschkow et al., 2021; Somai et al., 2020). These findings suggest that the brain uses its limited representational resources very sparsely, supporting an energy-efficient theory of working memory (Somai et al., 2020; Van der Stigchel, 2020). This account predicts that only the minimal amount of information is stored to the largest effect.

Importantly the minimal amount of information that we decide to store internally in order to perform a certain task, was shown to be dependent on a multitude of environmental factors such as locomotive demands (Draschkow et al., 2021) and time (Somai et al., 2020). When large movements are required between the encoding and retrieval of single objects in working memory, participants tend to encode more items in between movements, effectively optimizing the relative energy costs of movement and storage (Draschkow et al., 2021). While these findings nicely demonstrate the behavioral outcomes of this tradeoff, they do not reveal if similarly efficient resource allocation can occur for the representation of different object features. If only a specific feature of an object is relevant for the current task, are representational resources only used to represent that feature or all features constituting an object? In other words: Does the relevance of an object’s feature determine how well that feature is represented in the brain? In this study we aimed to investigate this question by manipulating the relevance of object features and quantifying the multivariate evidence on that feature in the EEG signal.

In addition, we set out to explore if relevance similarly effects the strength of neural representations when features are memorized as compared to when they are currently perceived. In contrast to working memory which represents information that is no longer available, we can define perception as a representational mechanism for information that is still available. Traditionally perception has been viewed as nearly unlimited in its capacity, an intuition that was doubtlessly influenced by our incredibly rich subjective experience of the world. But as we have established earlier, representations cost resources and converging evidence suggests that perceptual and working memory representations are both equally limited. In a recent study the authors presented participants with a classical change detection task in which several colored objects were either presented shortly (memory or absent condition) or for an extended period of time (perception or present condition; Tsubomi et al., 2013). Subsequently an identical number of colored stimuli was presented and participants had to indicate if any of the objects had changed color. Importantly in the present condition there was no empty delay interval between memory and probe and hence in this condition working memory should not be engaged in a traditional way. The authors reported that performance in both conditions was identical, suggesting that perception and working memory have similar capacity limitations. Furthermore, they showed that the contralateral delay activity, a common electrophysiological marker for working memory load was indistinguishable between both conditions.

The current study aims to answer several questions. First, we want to test if the relevance of an objects feature influences the strength with which this feature is represented in the EEG signal. Second, we want to test if the perceptual availability of a feature influences the strength with which this feature is represented in the EEG signal. We investigate this in a blocked 2 by 2 experimental designs in which participants were presented with colored gratings. These gratings were either presented for 150 ms (absent condition) and hence had to be remembered or remained on the screen for an extended period of time (1850). In half of the trials participants were asked to compare the gratings color to a subsequent probe, in the other half participants had to compare the spatial frequency. We used time-resolved multivariate pattern analysis to decode the spatial frequency of the stimuli in all four conditions. It was recently demonstrated that decodable, working memory related signals in the EEG can be explained by systematic eye movements and do therefore not necessarily originate directly from neural sources. This was primarily shown for the decoding of spatial orientation features that can trigger eye-movements to specific locations on the screen, even when stimuli are maintained in working memory (Mostert et al., 2018). Therefore, a gaze position based decoding analysis was conducted to control for the potential effects of eye-movements on the EEG decoding.

Our findings indicate that both relevance and availability influenced the multivariate evidence for the stimulus spatial frequency. Whereas high relevance reliably increased the amount of multivariate evidence, availability instead decreased the amount of multivariate evidence on spatial frequency.

These findings are particularly remarkable as stimulus features in the present condition were directly perceived and hence appeared far more vivid as compared to when the same feature was maintained in working memory.

## Materials and Methods

### 1. Participants

26 participants (aged 21-30, 16 females) with normal or corrected to normal vision enrolled in the experiment. None of the participants reported a history of psychiatric diagnosis. Informed consent forms were signed before the experiment and the study was approved by the local ethics committee. Subjects were compensated with 10 Euro/hour.

### 2. Stimuli

Stimuli consisted of vertical gratings (diameter 10° dva) with spatial frequency randomly selected from a set of 48 frequencies on every trial (1 cpdva to 4 cpdva, equally spaced) and were presented at fixation. Stimulus color was selected randomly from a set of 48 colors drawn from a circle in CIELAB color space (L = 54, a = 18, b = -8, radius = 59). A circular patch around the fixation point with radius 0.5° dva was cut out and the inner edges of the resulting annulus were blurred in order to prevent participants from basing their spatial frequency judgement on the relative position of grating and fixation cross. Similarly, we blurred the outer edges of the gratings since spatial frequency judgements might be based on the patterns of these sharp transitions zones. Stimuli were presented on an LCD display (27-inch, 2560 × 1440 resolution, 120 Hz refresh rate) using the Psychophysics Toolbox running in MATLAB (MathWorks). Participants were seated 58 cm away from the screen on a chinrest to prevent excessive head movements.

### 3. Protocol

Participants performed a blocked delayed match-to-sample task (Figure 1). Participants were instructed to remember the spatial frequency or the color of vertically oriented gratings and compare them to the respective feature of the subsequent probe. At the beginning of each block participants were informed about the relevant feature in this particular block (“Attend Color” versus “Attend Spatial Frequency”). Furthermore, participants were informed if the sample would stay on the screen or be removed during the delay interval (“Present Condition” versus “Absent Condition”). In the present condition the sample was presented for a total duration of 1850 ms followed by an empty screen for 150 ms. In the absent condition the sample was presented for 150 ms followed by an empty screen for 1850 ms. The relevance and presence were kept constant for the entire block. Hence participants had clear expectations on how long the stimuli would be available. 500 ms after the onset of the probe the fixation cross turned green, instructing the participant to report if the relevant feature was the same or different (50% probability) as compared to the sample. Participants performed a total of 1000 trials. The number of trials per condition was matched. To keep the task equally difficult between all conditions we used an online staircase procedure (Psychtoolbox QUEST algorithm) that was updating a psychometric function based on performance on every trial. The difference in spatial frequency and color between sample and probe was chosen based on this dynamic psychometric function so that performance was approximately at 75% in all conditions (1. Attend color absent 2. Attend color present 3. Attend SF absent 4. Attend SF present). At the end of every block participant received feedback in the form of a percent correct report. For practice all participants performed 5 trials of each condition at the beginning of the experiment where immediate feedback was provided.

**Figure 1.**
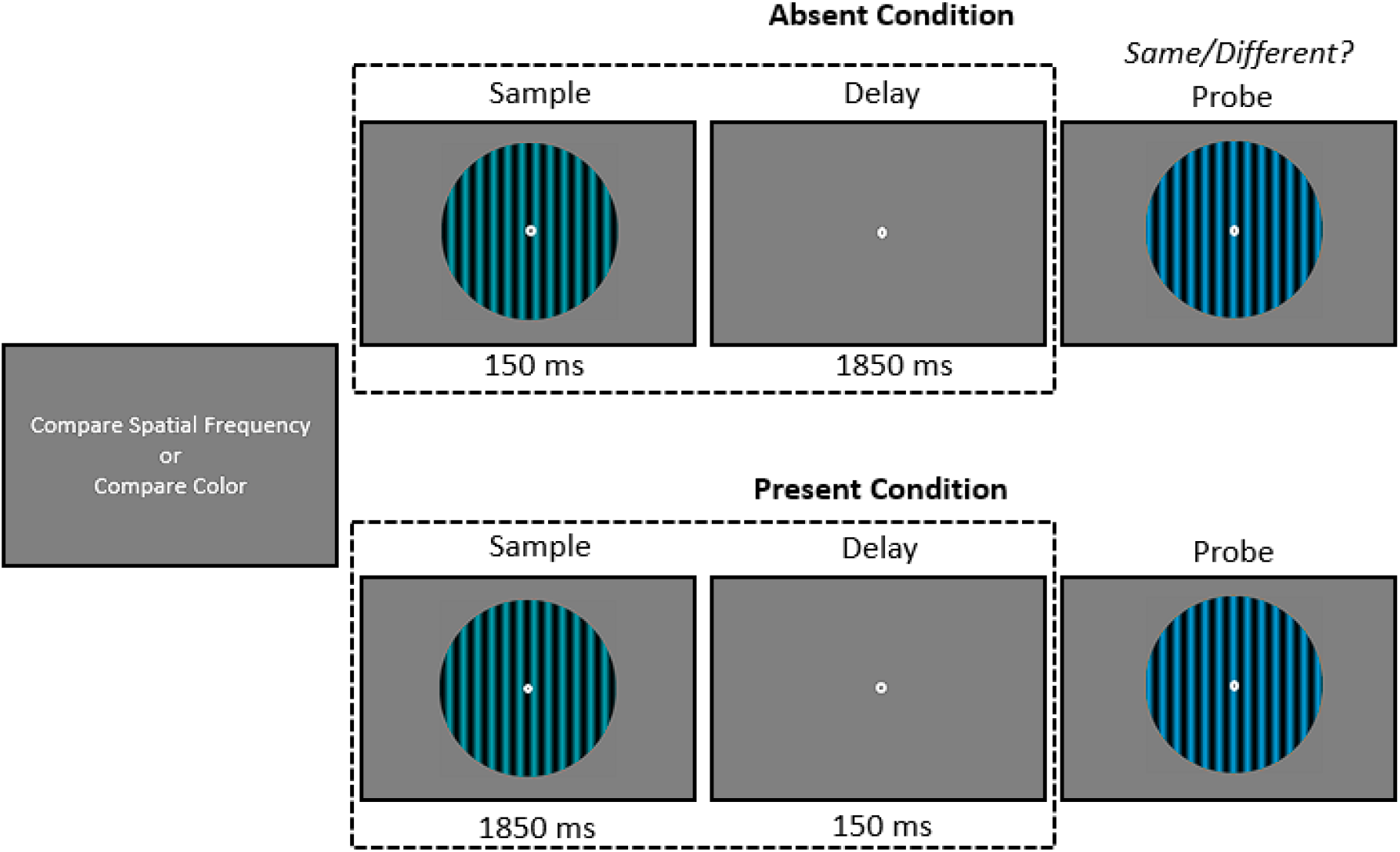
[Experimental Paradigm. Participants performed a delayed match-to-sample task. In ‘‘compare spatial frequency’’ blocks participants reported if the spatial frequency of the probe different from the sample. Similarly, in ‘‘compare color’’ blocks probe color was compared to samples. The duration for which the sample was presented on the screen was also varied in a blocked fashion, leading to 4 different types of blocks (compare SF absent, compare SF present, compare Color absent, compare Color present).]

**Figure 2.**
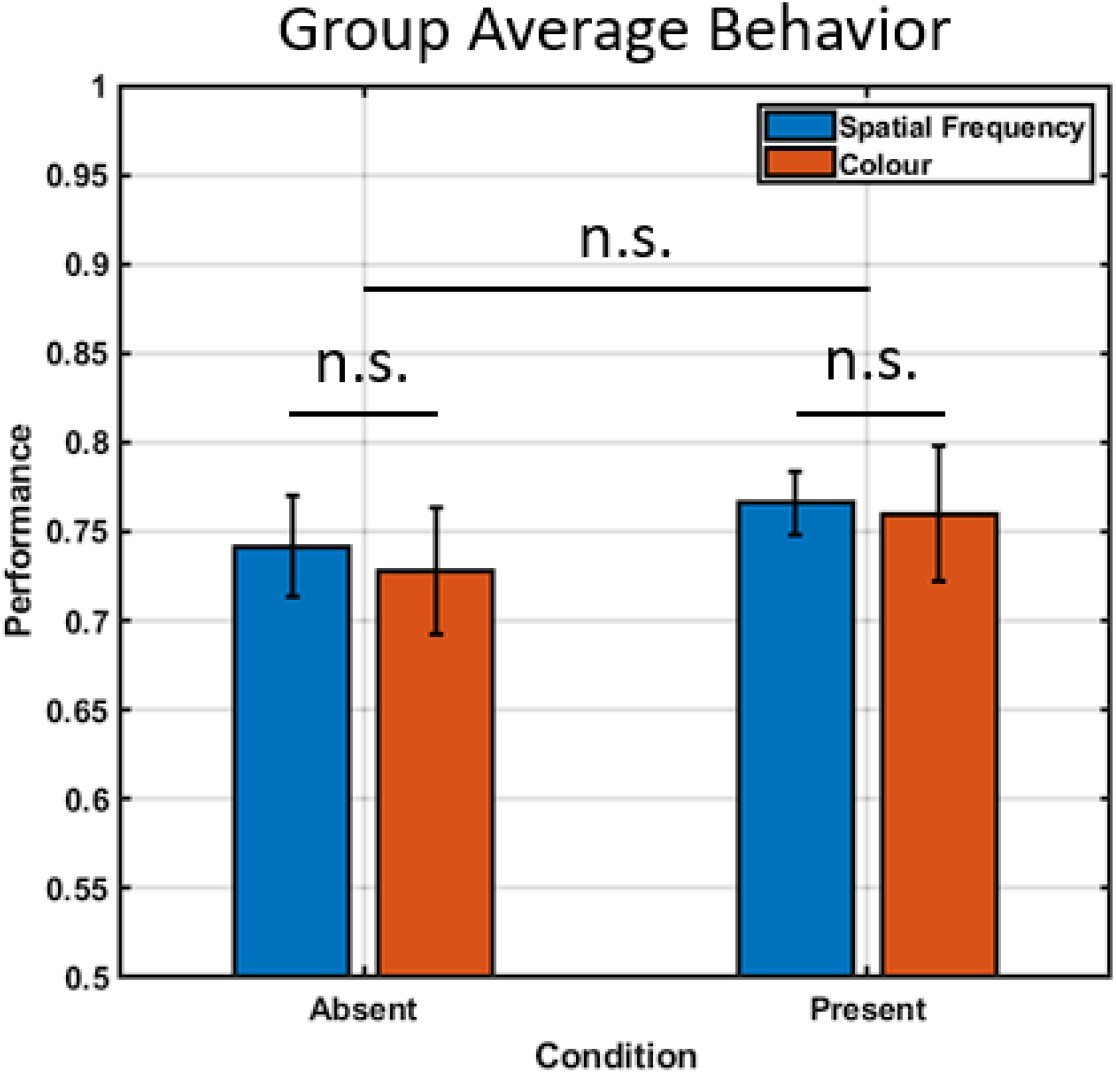
[Behavioral Accuracy. Group average percentage correct for all 4 conditions. Errorbars indicate standard error of mean.]

### 4. Eye Tracking Recording and Analysis

Gaze position was continuously tracked using an Eyelink 1000 (SR Researcher) eye tracker. Eye tracker calibration was performed at the beginning of the experiment and the gaze position was sampled at 1000 Hz. Saccades were extracted directly from the EyeLink saccade detection algorithm. Per condition we calculated the average number of saccades per trial as well as the average saccade amplitude.

For the Gaze position decoding analysis, we separated trials (vertical and horizontal position over time) into 4 different groups corresponding to the spatial frequencies used in the EEG decoding procedure. We performed baseline correction (−500 ms to 0 ms relative to stimulus onset) on the signal to remove slow drifts. Similar to the EEG decoding analysis we divided the gaze trials into three equal groups of which two served for training and one served as test set (3-fold cross-validation). MATLAB’s *fitcecoc* function (SVM, one-versus-all) was used to fit linear classifiers on two of the folds and test it on the third fold. Training and testing were performed on the same time point and was repeated 10 times, each time shuffling the trials that were divided into the three folds.

### 5. EEG recording and preprocessing

We recorded participants EEG using a 64 channel ActiveTwo Biosemi system. Two additional electrodes placed on the outer eye canthi recorded horizontal and vertical eye movements. Data analysis was performed in Matlab using the Fieldtrip toolbox. Prior to all preprocessing steps we identified and removed bad channels via visual inspection. The EEG data were then re-referenced to the average of channel T7 and T8, bandpass filtered between 0.01 and 80 Hz and line noise was removed using a DFT-filter (50 Hz). Thereafter the data were epoched from 1.5 s before sample onset to 3.5 s after sample onset. Large muscle and head-movement related artifacts were first removed through visual inspection. Afterwards we performed an ICA on the datasets separately to remove eye-movement related artifacts. Finally, the data were down-sampled to 100 Hz and absolute baseline correction was performed (window -500 ms to 0 ms before sample onset). We removed two subjects from the analysis due to a large number of EEG artifacts (>25% trials removed) caused by excessive head or eye movements during the experiment.

### 6. Multivariate Pattern Analysis

We decoded the spatial frequency of the sample stimulus based on the scalp topographies of the ERP voltage using a similar procedure as Bae & Luck (2018). We chose to train our decoders only on trials in which participants provided correct responses as these trials would reflect proper attention and memory allocation. Single trial ERP’s were lowpass filtered using an infinite impulse response butterworth lowpass filter (cutoff frequency 8 Hz) and subsequently down-sampled to 50 Hz. We decided to decode the spatial frequency from a selection of 20 occipital channels (PO7, PO3, O1, Oz, POz, Iz, PO8, PO4, O2, P1, P2, P3, P4, P5, P6, P7, P8, P9, P10, Pz) as we were mainly interested in visual representations.

For every participant we binned trials into 4 spatial frequency classes of equal size. Trials in every class were divided into 3 equally sized groups and averaged, resulting in 3 folds that were later used for 3-fold cross-validation. We used MATLAB’s *fitcecoc* function to fit multi-class error correcting output code models using support vector machines that were trained to distinguish between one class and all others (one-versus-all). For every individual separately, classifiers were trained on single timepoints and received two samples from each of 4 SF classes consisting of 20 features in the form of EEG topography voltage (20 channels). Classifiers were then tested on all timepoints using the remaining fold (4 classes, 20 features). This procedure was repeated 3 times with each fold serving as training and test set once, but never as both simultaneously. In addition, we randomly shuffled the trials that were averaged into each fold 10 times, each time repeating the entire decoding procedure. The classifiers output scores (distance-to-bound) were z-scored, scores corresponding to samples that were not present on that trial were multiplied with -1 and all scores were averaged.

### 7. Statistical Analysis

We assessed statistical significance of our decoding accuracy time-series using a nonparametric cluster-based permutation test (Maris & Oostenveld, 2007). This was done by first simulating the performance of our decoder that would be observed when guessing randomly. For every participant and timepoint we randomly generated either a correct or a false classification value by multiplying the distance-to-bound scores with 1 or -1. We then ran t-tests for every timepoint of the group-averaged decoding time-series and identified clusters of consecutive significant tests. The t-values within each cluster were summed to calculate the cluster-level t mass. This procedure was repeated times to generate a distribution of cluster-level t masses. We then compared the actually observed cluster-level t masses to the null-distribution and tested if their mass exceeded the 95% quantile. Clusters were considered significant if they were larger than 95% of simulated clusters. The same procedure was performed to estimate clusters of significant above chance decoding for the gaze decoding.

To compare 1-D cluster sizes between conditions we performed a cluster-based permutation test by randomly multiplying the decoding scores of individual validation runs by -1 and recalculating the difference in significant cluster sizes between conditions. This was performed 20.000 times to generate the null-distribution under the null hypothesis (H0: EEG trials do not contain information regarding the spatial frequency). The veridical difference in significant cluster size was then compared to the null-distribution and was considered significant ifs value exceeded the 95% quantile of the null-distribution.

Number of saccades and saccade amplitude was statistically analyzed using two-way ANOVAs with factors availability (absent, present) and relevance (spatial frequency, color) as well as students t-tests.

## Results

### 1. Behavior

A One-Way ANOVA with factors relevance (SF vs Color) and availability (absent vs present) revealed no main effect of relevance (F(1,108) = 1.683, p = 0.197) or availability (F(1,108) = 1.483, p = 0.226) and no interaction effect between the two (F(1,108) = 0.037, p = 0.847), indicating that our staircase procedure successfully equalized task difficulty between conditions.

### 2. Effect of Availability and Relevance

We trained linear classifiers to quantify how much evidence for the spatial frequency of a stimulus was present in the EEG signal over time. Stimuli were presented either for 150 ms followed by a blank delay interval of 1850 ms (absent condition) or remained on the screen for 1850 ms, followed by a blank delay interval of 150 ms. In the compare spatial frequency condition participants had to indicate if the probe had a similar or different SF as compared to the sample. In the color condition participants indicated if the stimulus color was the same or different between sample and probe.

Visual inspection of the MVPA temporal generalization matrices indicated that both relevance and availability modulated decodability (Figure 3ABDE). Statistical analysis of the timepoint-by-timepoint decodability revealed that SF could be decoded longer if it was the relevant feature as compared to when it was irrelevant in both absent and present conditions (Figure 3 G: *absent relevant 1180 ms, absent irrelevant 380 ms*, cluster-based permutation test: p<0.0125, Figure 3 H: *present relevant 660 ms, present irrelevant 420 ms*, cluster-based permutation test: p<0.0125). Furthermore, SF could be decoded more accurately if it was relevant in the absent condition as compared to when it was irrelevant, although this was not continuously the case (700 ms to 1160 ms, Figure 3E).

**Figure 3.**
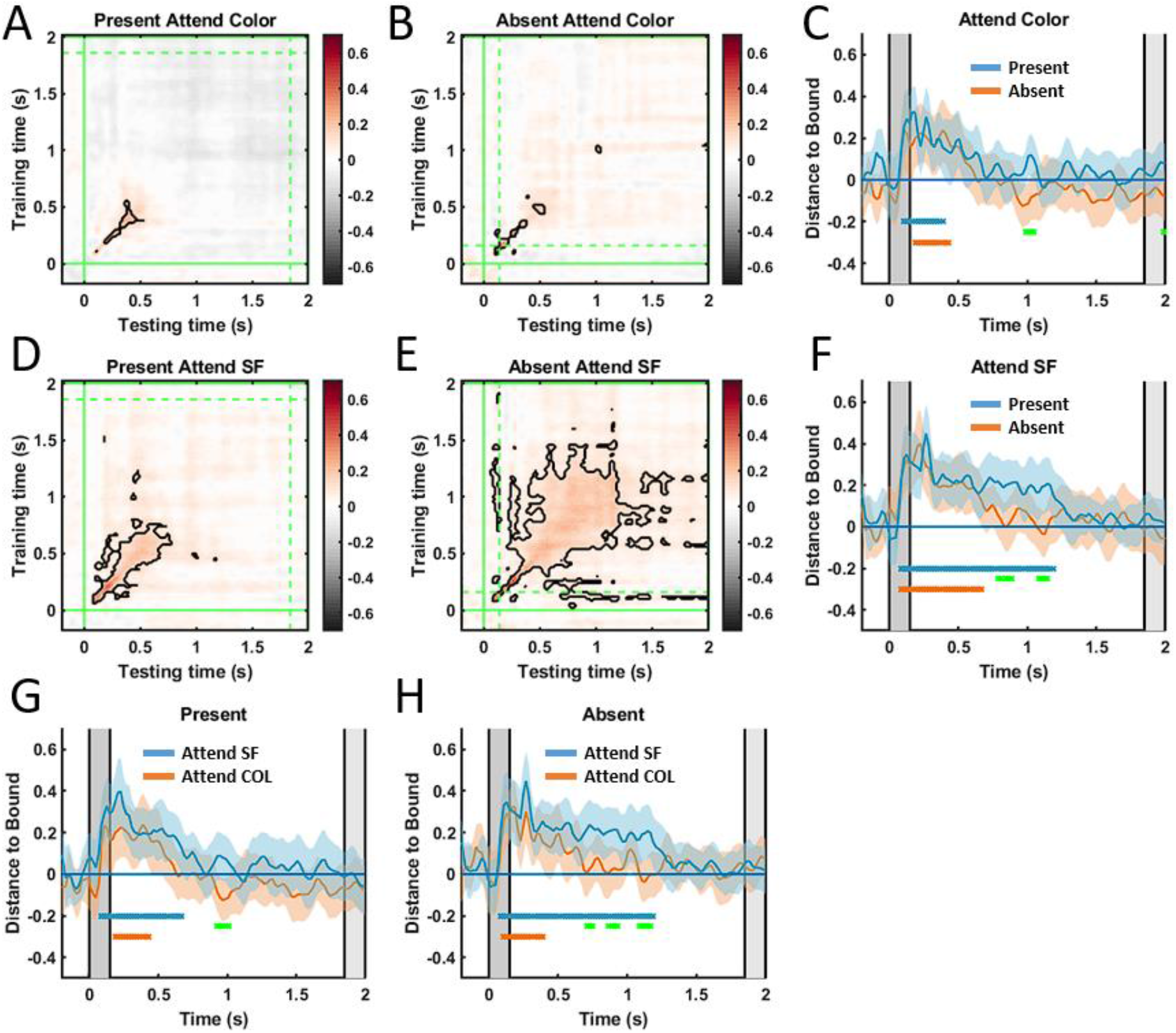
[EEG Decoding. **A**. Temporal generalization decodability matrix for Present Attend SF condition. Y axis indicates timepoints on which classifiers were trained. X axis indicates timepoints on which classifiers were tested. Color axis indicates mean normalized distance-to-bound scores. Green lines indicate onset (solid) and offset (dashed) of stimuli in the respective condition. **B**. Same but for Absent attend color, **D** Present attend SF and **E** Absent attend SF conditions. **C**. Diagonal decoding performance (timepoint-by-timepoint) of absent (blue line) and present (orange line) attend color conditions. Colored areas indicate 95% confidence intervals. Colored dots (blue, orange) indicate significant clusters of above chance decoding. Green dots indicate significant differences between absent and present condition (t-tests). Gray bars in the background indicate onset and offset of stimulus in absent and present conditions respectively. **F**. same but for attend SF absent (blue) and present (orange) conditions, **G**, present attend SF (blue) and attend color (orange) conditions, **H**, absent attend SF (blue) and attend color (orange) conditions.]

Most importantly we found that SF could be decoded longer in the absent condition as compared to the present condition when SF was relevant (absent: 1180 ms, present 660 ms, cluster-based permutation test: p<0.0125, Figure 3 F). This result is particularly striking because participants were directly fixation and perceiving the grating in the present condition for almost 2 seconds and were aware that the spatial frequency was relevant for successful probe comparison. Significant decoding clusters did not differ in length between absent and present conditions when SF was irrelevant (cluster-based permutation test: p>0.1875, Figure 3 ABC). This indicates that the larger decoding cluster found in the absent SF relevant condition was not caused by the offset of the sample stimulus at time 150 ms.

We hypothesize that the lack of sustained decoding in the present relevant condition might be a result of the long availability of the stimulus. As participants were aware that the relevant feature will be perceptually available in the external world for at least 1850 ms, the visual system likely has no incentive to engage large amounts of resources to represent the feature. Alternatively, participants might have not attended the stimulus in the present condition as much as the only shortly available stimulus in the present condition, leading to differences in decoding performance. Our following analysis aimed to address this possibility.

### 3. Attention Decoding

To test whether participants dedicated few or none of their attentional resources to the present stimulus at its onset we conducted an additional decoding analysis. Our rationale was as follows: If participants did not attend the present stimulus at its onset (∼0 to 500 ms) because they were aware of its long availability then the neural signals during that time in present relevant and present irrelevant conditions should not be distinguishable. Conversely, if participants attended the present stimulus, then a classifier should be able to distinguish between present relevant and irrelevant conditions.

We trained a new set of binary support vector machines to distinguish between trials from the present relevant and irrelevant conditions, regardless of their spatial frequencies. We find that the classifier can distinguish between present relevance conditions as early as 187 ms after stimulus onset indicating that attentional signals differed between them (Figure 4AC). This finding, together with the fact that we found a significant effect of relevance in the present condition, suggests that the poor SF decoding found in the present relevant condition was likely not caused by a lack of attention at stimulus onset.

**Figure 4.**
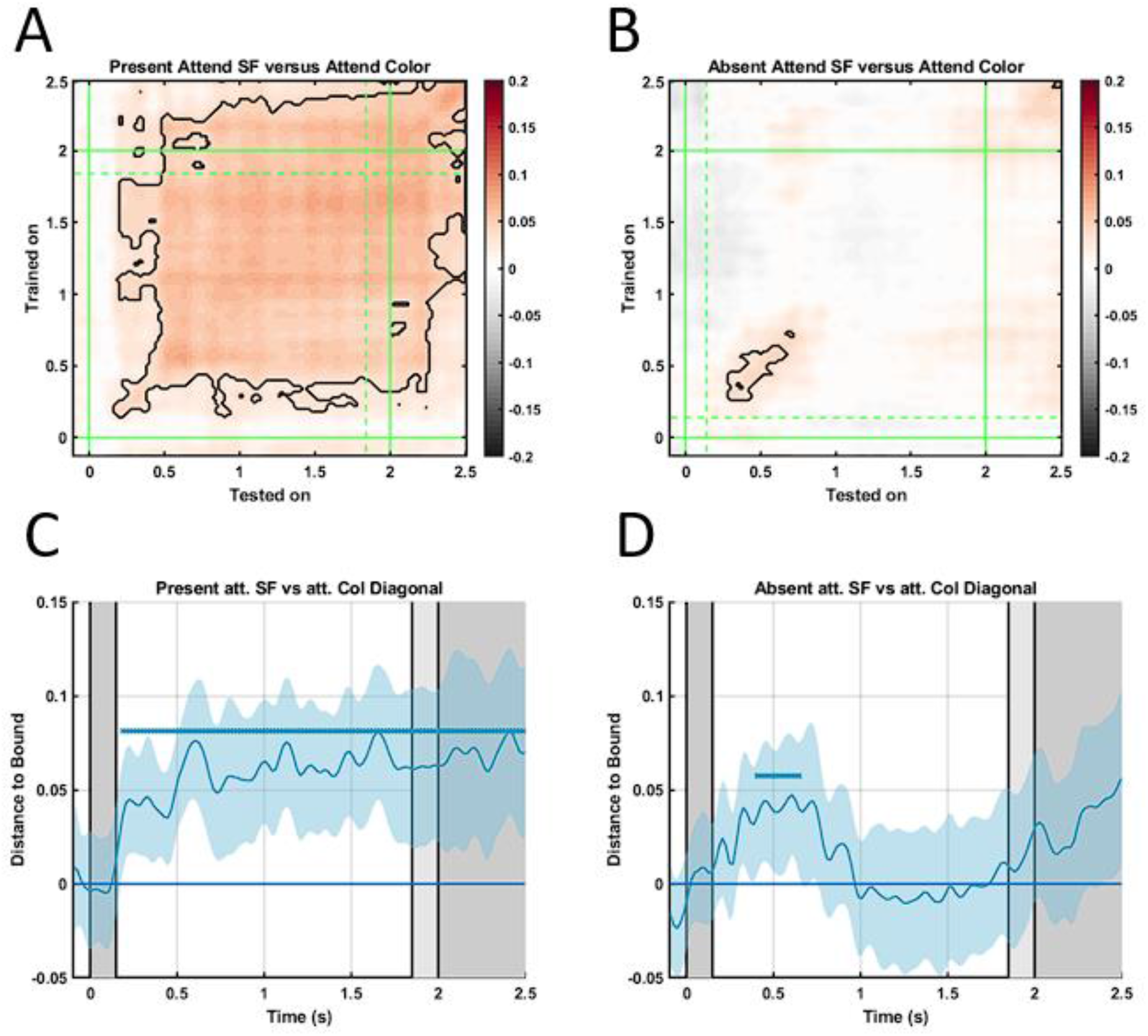
[Attention EEG Decoding. **A**. Temporal generalization decodability matrix for SF relevant versus irrelevant. Color axis indicates mean normalized distance-to-bound scores. Green lines indicate onset (solid) and offset (dashed) of stimuli in the respective condition. **B**. Same but for absent condition. **C, D**. Diagonal decoding (timepoint-by-timepoint) of and present and absent conditions. Colored areas indicate 95% confidence intervals. Blue colored dots indicate significant clusters of above chance decoding.]

### 4. Saccade Analysis

Eye movements were previously shown to systematically track orientation features held in working memory and can lead to decodable artifacts in the EEG signal (Mostert et al., 2018). We considered this possibility in the design of our task and choose to use spatial frequency instead of spatial orientation as decodable feature, as this would not provide spatial locations, such as orientation endpoints to bias eye movements that were correlated with our decoding variables. To test if this was successful, we attempted to decode the SF from gaze position alone. Multi-class classifiers could not distinguish between different SF based on gaze location (horizontal, vertical) in any condition (Figure 5). Our EEG decoding results are therefore unlikely to be a result of eye-artifacts caused by systematic eye movements.

**Figure 5.**
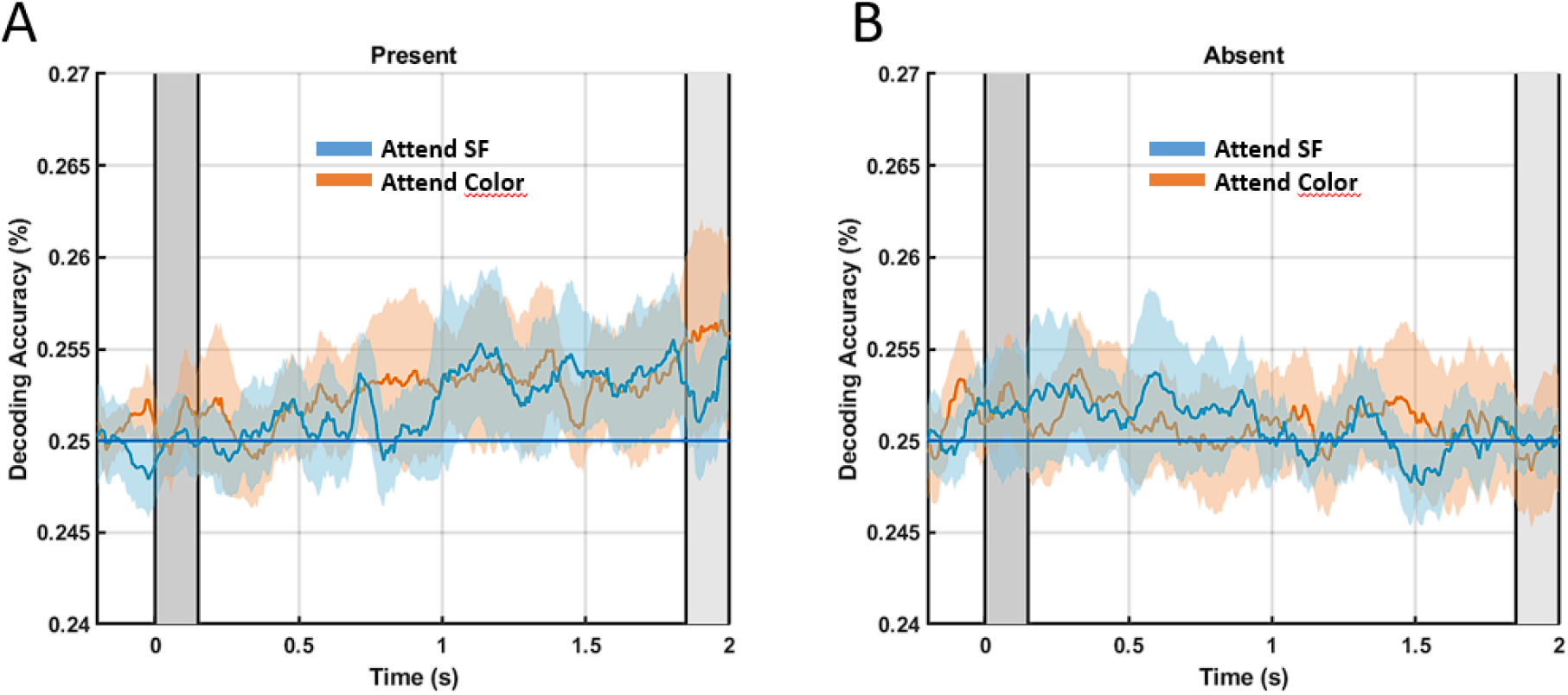
[Saccade Decoding SF. Timepoint-by-timepoint decoding of SF from Gaze positions. **A**. Present condition SF relevant (blue) and irrelevant (orange). Colored areas indicate 95% confidence intervals. Gray bars in the background indicate onset and offset of stimulus in absent and present conditions respectively. Colored dots (blue: SF relevant, orange: SF irrelevant) indicate significant clusters of above chance decoding. **B**. same but for Absent condition.]

## Discussion

In this study we tested how two task factors, feature availability and feature relevance, influence the amount of multivariate evidence found for that feature in the EEG signal. We hypothesized that high feature relevance would lead to more multivariate evidence, which was confirmed by our decoding analysis. Furthermore, we hypothesized that long availability of a stimulus feature would lead to less multivariate evidence which we could also confirm using MVPA. Our findings suggest that the visual system efficiently evaluates how much resources to dedicate to the representation of stimuli and that factors such as time of perceptual availability and relevance bias this evaluation.

How can our findings be explained in the light of commonly observed reduction in visual activity over time, usually referred to as visual adaptation (Webster, 2015)? Visual adaptation has been observed on multiple time-scales in the visual system and can occur in time ranges matching the average duration of a fixation (∼400 ms) (Akyuz et al., 2020; Gutnisky & Dragoi, 2008). Although visual adaptation likely plays an important role in explaining the reduced decodability found in the present condition there still remain several open questions. First and foremost, how does the brain generate vivid visual experience if activity in visual areas is reduced as a result of adaptation? Evidently our decoding analysis might simply not be sensitive enough to reveal the signals generating subjective experience. Simultaneously however we found highly significant decodable signals in the absent condition when the stimulus is presumably in working memory and less vividly perceived. In this particular setup we hence find multivariate evidence to be anticorrelated with subjective experience.

This apparent contradiction could be explained in several ways. Although intuitively one might assume that experience should correlate with neural evidence this does not necessarily have to be true. On the one hand this assumption is fundamentally dependent on our definition of evidence (e.g., precision, signal to noise) and on the other hand the brain might be able to generate vivid subjective experience from relatively sparse activity. In the latter case the question arises of what the increased decodability of the feature in the absent condition actually reflects. It is possible that it indicates actual representational effort related to the maintenance of the stimulus in working memory. This additional effort might not be necessary in the present condition where the stimulus is ‘‘automatically’’ represented in the forward stream of the visual system. Another possibility is that the stimulus in the present condition is simply represented less strongly in neural signals in the present condition since participants were fully aware that they will not be needing precise representations of the particular feature in the early parts of the trial. This possibility might lead to the intriguing conclusion that subjective experience does not actually reflect how well a visual feature is represented internally.

Our findings can also be interpreted in the context of predictive coding (Friston & Kiebel, 2009). Hierarchical predictive coding theories of visual processing state that hierarchically higher areas of the visual pipeline continuously make predictions to explain the activity generated in hierarchically lower areas of the visual processing pipeline. In this framework only information that does not match the predicted responses, the so-called prediction error, is forwarded higher into the hierarchy whereas correctly predicted signals are dampened. It is not entirely clear if this process leads to increases or decreases in the decodability of perceived features, as some evidence suggests that similar representations are sharpened and dissimilar representations are suppressed (Kok et al., 2012; Panichello et al., 2013). Also how this translates to the increased decodability of SF in the absent condition remains a highly interesting question that future work will have to address (Parr & Friston, 2017).

Although our MVPA analysis revealed no evidence at all for the spatial frequency after a few hundred of milliseconds (absent: 1180 ms, present 660 ms) it is unlikely that the information completely vanished from the brain and more likely reflects the limitations of our EEG MVPA approach. As participants were able to perform the task with reasonable accuracy the information must have been somehow stored but escaped our classifiers. This could be due to a number of reasons. First it is plausible that the quality of our signal deteriorated over time due to slow drifts in the EEG signal (van Driel et al., 2021). It has also been proposed that neural representations can be stored in so called ‘‘activity-silent’’ states that rely on short-term synaptic changes to encode information (Chota & Van der Stigchel, 2021; Mongillo et al., 2008). This latter possibility has mostly been discussed in the context of working memory; hence it is not clear if this also applies to the representation of perceptual information.

Functionally the absence of decodable information in the present condition could also be explained by the fact that participants did not have to explicitly represent the exact spatial frequency. As the empty delay between stimulus and probe only lasted 150 ms, participants might have been able to use sensory perceptual traces such as iconic memory to detect changes. Importantly this strategy would not require participants to have an internal representation of the feature at all since the change would immediately pop out. We tried to minimize the effectiveness of this strategy by shifting the phase of the probe’s spatial frequency on every trial and hence create additional transients to cover up the changes in SF. Potentially this might have forced participants to attend spatial frequency channels in particular, leading to the observed differences in SF relevant and SF irrelevant conditions. In future work we aim to investigate this by increasing the delay between stimulus and probe while still keeping stimuli perceptually available for an extended period of time.

## Conclusion

In conclusion we found that high feature relevance led to increased multivariate evidence for perceived and memorized stimuli. Furthermore, we also showed that stimulus features are better represented when they are perceptually available for a short as compared to a long duration. Our findings outline a number of factors that determine how well we represent visual information internally and raise the intriguing question of why multivariate evidence from visual areas is not indicative of subjectively experience vividness of visual features.

